# Molecular determinants for α-tubulin methylation by SETD2

**DOI:** 10.1101/2020.10.21.349365

**Authors:** Sarah Kearns, Frank M. Mason, W. Kimryn Rathmell, In Young Park, Cheryl Walker, Kristen Verhey, Michael A. Cianfrocco

## Abstract

Post-translational modifications to tubulin are important for many microtubule-based functions inside cells. A recently identified tubulin modification, methylation, occurs on mitotic spindle microtubules during cell division, and is enzymatically added to tubulin by the histone methyltransferase SETD2. We used a truncated version of human SETD2 (tSETD2) containing the catalytic SET and C-terminal Set2 Rpb1 interacting (SRI) domains to investigate the biochemical mechanism of tubulin methylation. We found that recombinant tSETD2 has a higher activity towards tubulin dimers than polymerized microtubules. Using recombinant single-isotype tubulin, we demonstrate that methylation is restricted to lysine 40 (K40) of α-tubulin. We then introduced pathogenic mutations into tSETD2 to probe the recognition of histone and tubulin substrates. A mutation in the catalytic domain, R1625C, bound to tubulin but could not methylate it whereas a mutation in the SRI domain, R2510H, caused loss of both tubulin binding and methylation. We thus further probed a role for the SRI domain in substrate binding and found that mutations within this region had differential effects on the ability of tSETD2 to bind to tubulin versus RNA Polymerase II substrates, suggesting distinct mechanisms for tubulin and histone methylation by SETD2. Lastly, we found that substrate recognition also requires the negatively-charged C-terminal tail of α-tubulin. Together, this work provides a framework for understanding how SETD2 serves as a dual methyltransferase for histone and tubulin methylation.

## Introduction

Microtubules are dynamic cytoskeletal polymers that maintain cell shape, serve as tracks for intracellular trafficking, provide a structural framework for cell division, and form the structural elements of cilia. How microtubules achieve their many varied cellular functions comes, in part, from a tubulin code of multiple isoforms of α- and β-tubulin dimers and varied post-translational modifications (PTMs) (1–3). For example, differentiated cells express varying amounts of α- and β-tubulin isotypes to perform specialized roles (4). Additionally, within each cell there are subpopulations of microtubules with PTMs that further regulate microtubule-based functions. Analogous to how the histone code directs chromatin function, the combination of tubulin isotypes and PTMs comprise a tubulin code that specializes microtubule function in cells.

One key chromatin modifier that contributes to both the histone and tubulin codes is SET domain containing 2 (SETD2). SETD2 is an S-adenosyl-methionine (SAM)-dependent lysine methyltransferase, which on chromatin is responsible for histone-3 lysine 36 trimethylation (H3K36me3), a mark associated with gene transcription (5, 6). Loss of SETD2 is embryonic lethal in part because its ability to trimethylate H3K36 is non-redundant (7). Many cancers including kidney, lung, bladder, glioma, and leukemia have inactivating mutations in SETD2 (8–12). In clear cell renal cell carcinoma (ccRCC), SETD2 is the second most frequent mutation contributing to 10-15% of all ccRCC cases (6, 13–16). For example, pathogenic arginine-to-cysteine mutation at position 1625 (R1625C), found within the catalytic SET domain, ablates methyltransferase activity and is associated with poor prognosis. Another mutation, arginine-to-histidine at position 2510 (R2510H), occurs in the Set2-Rpb1-interacting (SRI)-domain at the C-terminus of SETD2 but does not result in loss of H3K36me3 (17–19), suggesting that pathogenicity associated with this SRI domain mutation is not due to loss of histone methylation.

Our previous work demonstrated that SETD2 can methylate tubulin and that methylation occurs at Lysine 40 of α-tubulin (αTubK40me3) (19). In dividing cells, αTubK40me3 localizes to the minus ends of microtubules that form the mitotic spindle (19). Loss of SETD2 in ccRCC and knock-out of SETD2 in cells results in genomic instability and mitotic defects such as multipolar spindles, lagging chromosomes during anaphase, chromosomal bridging during cytokinesis, and micronuclei (19, 20). These phenotypes correlated with a drastic reduction in both H3K36me3 and αTubK40me3 methylation. Reintroduction of a truncated form of wild-type SETD2 (tSETD2) containing the SET and SRI domains rescued both histone and tubulin methylation as well as the mitotic defects (18, 19). In contrast, expression of tSETD2 with the R2510H mutation in the SRI domain rescued histone methylation but was unable to rescue tubulin methylation or the mitotic defects (18, 19), suggesting that a loss of SETD2 activity can result in increased of mitotic defects in a tubulin-dependent manner.

Many aspects of SETD2 function are still unexplained, such as how SETD2 recognizes and differentiates between methylation substrates. Here, we took a biochemical reconstitution approach to define the minimal components required to methylate tubulin *in vitro.* By utilizing both purified tSETD2 and recombinant human tubulin (21–23) we had precise control of tubulin isotype and PTM-state *in vitro* and allowed us to generate mutant versions to probe site selectivity of SETD2 methylation. We demonstrate that tSETD2 is sufficient to methylate tubulin *in vitro* and has a higher activity towards tubulin dimers over microtubules. We verify that the methylation site is on α-tubulin lysine 40 (αTubK40), the same site that can be acetylated (24–27). We also find that the SRI domain of SETD2 is important for binding tubulin substrate as the tSETD2-R2510H mutant can neither pull down nor methylate tubulin. Because the SRI domain of SETD2 makes protein-protein interactions with RNA Polymerase II (RNA Pol II) during transcription (28), we investigated residues that could distinguish binding between tubulin and RNA Pol II. We found that positively charged residues within the SRI domain are more important for tubulin binding where aromatic residues are more critical for RNA Pol II binding. In addition, we found that SETD2 recognizes the negatively charged C-terminal tail of α-tubulin, likely through electrostatic interactions. Together, this work further establishes SETD2 as a tubulin methyltransferase and provides a molecular basis for changes in tubulin and histone methylation derived from ccRCC mutations in SETD2 due to SRI-domain regulation.

## Results

SETD2 methylates tubulin *in vitro.* To reconstitute tubulin methylation *in vitro*, we purified a truncated form of SETD2 (tSETD2, aa1418-2564, Fig. 1A,B) containing both SET and SRI domains. The smaller size of tSETD2 made it more amenable for biochemical purification and previous work indicated that tSETD2 is sufficient to rescue SETD2 loss of function (18, 19). When expressed in mammalian cells, a FLAG-tagged version of tSETD2 (tSETD2-FLAG) localized to the nucleus during interphase and was dispersed throughout mitotic cells (Fig. S1–2), similar to the localization of expressed full-length and endogenous SETD2 (29, 30). Recombinant tSETD2-FLAG was purified from HEK293 cells, yielding a single species on size exclusion chromatography (Fig. 1C) and a single band detected by SDS-PAGE (Fig. 1D). We confirmed that this band was SETD2-FLAG via western blot against the FLAG epitope (Fig. 1D).

**Figure 1:**
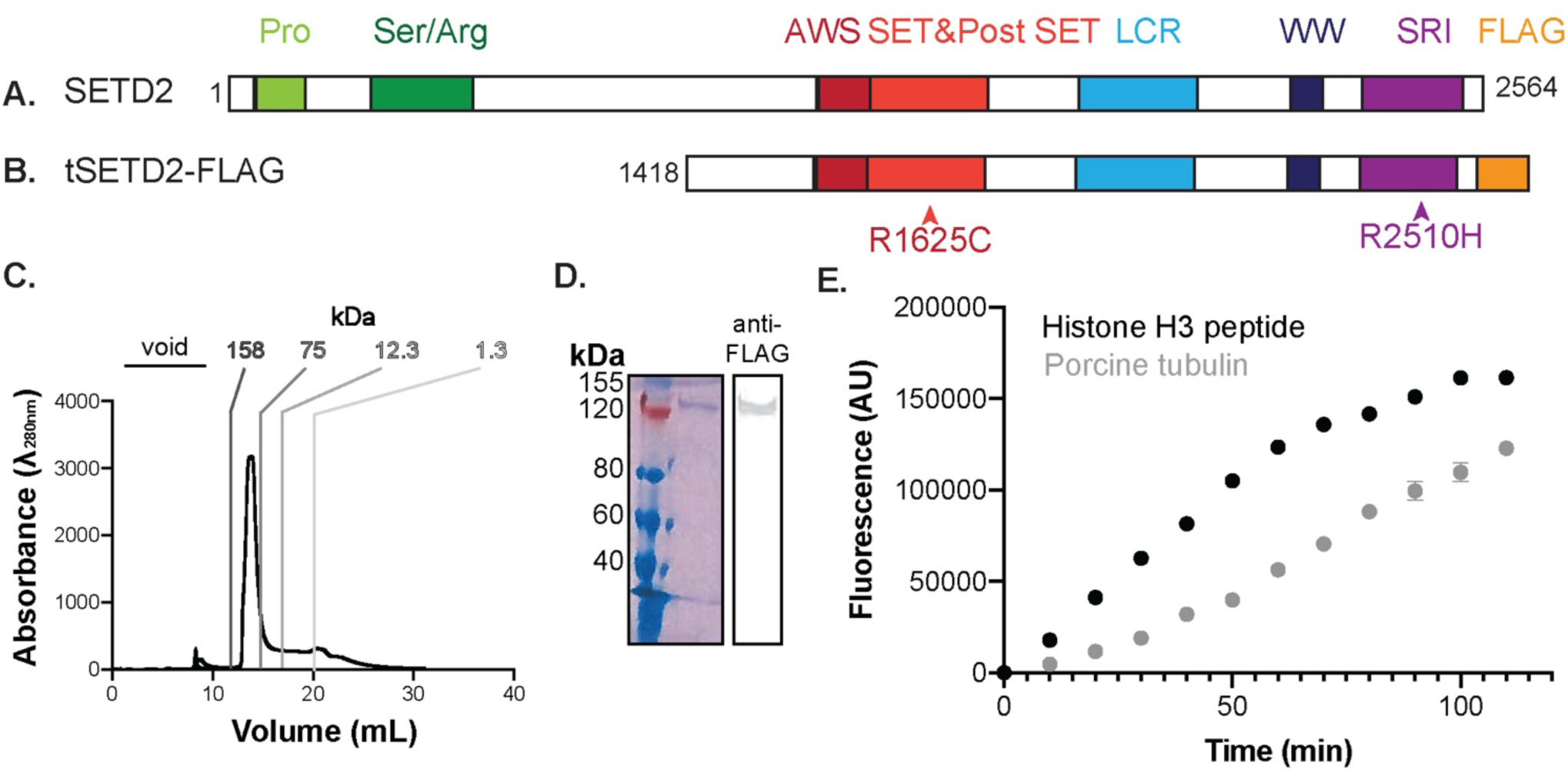
tSETD2-FLAG methylates tubulin *in vitro*. Schematic of A) full-length and B) tSETD2-FLAG domain organization. Pro: Proline-rich (light green), Ser/Arg: Serine/Arginine-rich (dark green), AWS: Associated with SET (dark red), SET (red), LCR: locus control region (light blue), WW: Tryptophan-rich (dark blue), SRI: Set2 Rpb1 interacting domain (purple). C-D) tSETD2-FLAG was purified from HEK293 cells and analyzed by C) size-exclusion chromatography and D) Coomassie-stained gel (left) and anti-FLAG western blot (right). E) Fluorescence-based assay of methyltransferase activity over time. tSETD2-FLAG was incubated with methyl donor SAM and 5 μM of either histone H3 peptide (aa 21-44, black) or porcine brain tubulin (grey) substrate. Data are presented as average±SD of n=4 experiments with two different tSETD2-FLAG purifications.

To measure the enzymatic activity of tSETD2-FLAG, we adopted a fluorescence-based assay that monitors S-adenosyl homocysteine production, the product of SAM-dependent methyl transfer. In this assay, an increase in fluorescence directly corresponds to tSETD2 activity. Thus, tSETD2-FLAG was incubated with porcine brain tubulin protein and the methyl donor SAM. As a positive control, we used an H3 peptide (residues 21-44) which can be methylated by tSETD2 (31). As a negative control, we measured fluorescence activity of tSETD2-FLAG in the presence of SAM but the absence of any substrate. In this case, any methyltransferase activity is indicative of auto-methylation that can then be subtracted out of substrate-containing reactions. Both histone peptide and tubulin protein substrates produced an increase in fluorescence activity over time, and tSETD2-FLAG activity was higher towards histone peptide than tubulin protein (Fig. 1E). K_M_ values for tubulin and H3 peptide substrates (0.75±0.06 μM and 0.45±0.32 μM, respectively) suggest that tSETD2-FLAG has slightly different affinities depending on substrate (Fig. S3). However, the V_max_ with tubulin protein as a substrate was lower compared to H3 peptide (Fig. S3), suggesting that the active site of SETD2 becomes saturated more quickly with tubulin than with H3 peptide. These data indicate that our reconstituted protein has activity towards tubulin substrate.

### tSETD2 displays a higher activity towards tubulin dimers than microtubule polymer

Our previous work showed that GST-SETD2(1392–2564) is capable of methylating tubulin in both soluble dimeric and polymerized microtubule states (19), however, the relative ability of SETD2 to methylate these substrates was not tested. To do this, we measured tSETD2-FLAG activity against microtubules maintained in a polymerized state with taxol as compared to porcine brain tubulin dimers maintained in an unpolymerized state with podophyllotoxin (32, 33) (Fig. 2A). We found that tSETD2-FLAG has higher activity toward tubulin dimers at 5 μM (5.08±0.06 nmol/min) than polymerized microtubules at the same concentration (3.19±0.05 nmol/min) (Fig. 2B-C). By varying the substrate concentrations, we generated Michaelis-Menten plots for both substrates. Calculation of the Michaelis constant, a measure of affinity, demonstrated that there is not a statistically significant difference in K_M_ for microtubule vs tubulin substrates (Fig. 2D). However, tSETD2-FLAG displayed a higher V_max_ for tubulin dimer (4.40±1.03 nmol/min) than microtubule (3.15±0.14 nmol/min) substrate (Fig. 2D). This finding is consistent with the K40 methylation site on α-tubulin, which is luminal in polymerized microtubules, being more accessible in a tubulin dimer context. Microtubule methylation may then occur via the incorporation of methylated dimers into microtubules.

**Figure 2:**
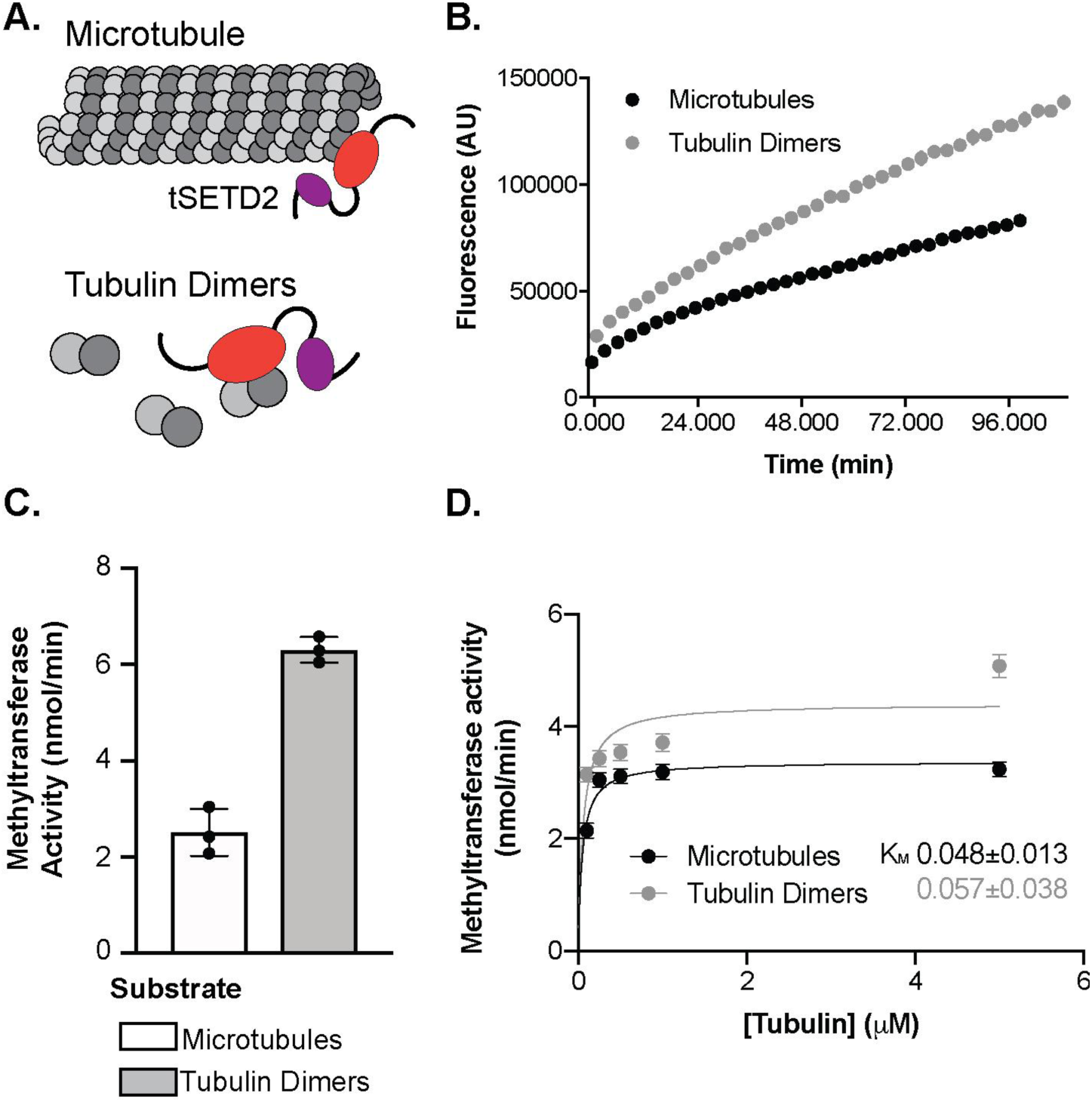
tSETD2-FLAG has higher activity towards tubulin dimers than microtubule polymer. A) Schematic showing tSETD2 (SET domain in red and SRI domain in purple) methylation on either tubulin dimers or polymerized microtubules, tubulin state ensured using small molecule podophyllotoxin or taxol, respectively. B-D) Purified tSETD2-FLAG was incubated with SAM and 5 μM porcine brain tubulin or microtubule substrates and methyltransferase activity was monitored over time. Shown are B) representative fluorescence over time traces and C) calculated average activity of tSETD2-FLAG with taxol-stabilized microtubules (white) or podophyllotoxin-maintained tubulin dimers (grey). Each dot indicates the average result from a single experiment and the bar represents the average±SD methyltransferase activity across n=3 experiments. D) Michaelis-Menten plot of the methyltransferase activity dependence on substrate concentration, either microtubules (black) or tubulin dimer (grey). Data are reported as average±SD methyltransferase activity from n=3 experiments.

### SETD2 methylates α-tubulin at lysine 40

We previously determined that SETD2 can methylate α-tubulin at lysine 40 (αTubK40) using recombinant GST-tagged SETD2(1392–2564) and porcine brain tubulin followed by mass spectrometry (19). However, brain tubulin contains numerous isotypes and pre-modified tubulin proteins and thus it has been difficult to determine whether there are additional sites for SETD2 methylation on α- or β-tubulin.

To determine if there are other methylation sites on tubulin, we utilized recombinant human tubulin. Recent advances in the expression and purification of recombinant single-isotype tubulin (21–23) allowed us to purify human αTub1B/βTub3 tubulin dimers (hereafter referred to as αβ-tubulin) from insect cells using the baculovirus expression system (Fig. 3A-B). This system allowed us to mutate the putative methylation site K40 on α-tubulin by purifying αβ-tubulin dimers with a mutation of K40 to alanine [αβ-tubulin(αK40A)]. We confirmed that αβ-tubulin, both WT and K40A, were functional by observing their ability to polymerize into microtubules from GMPCPP-tubulin seeds (Fig. S4).

**Figure 3:**
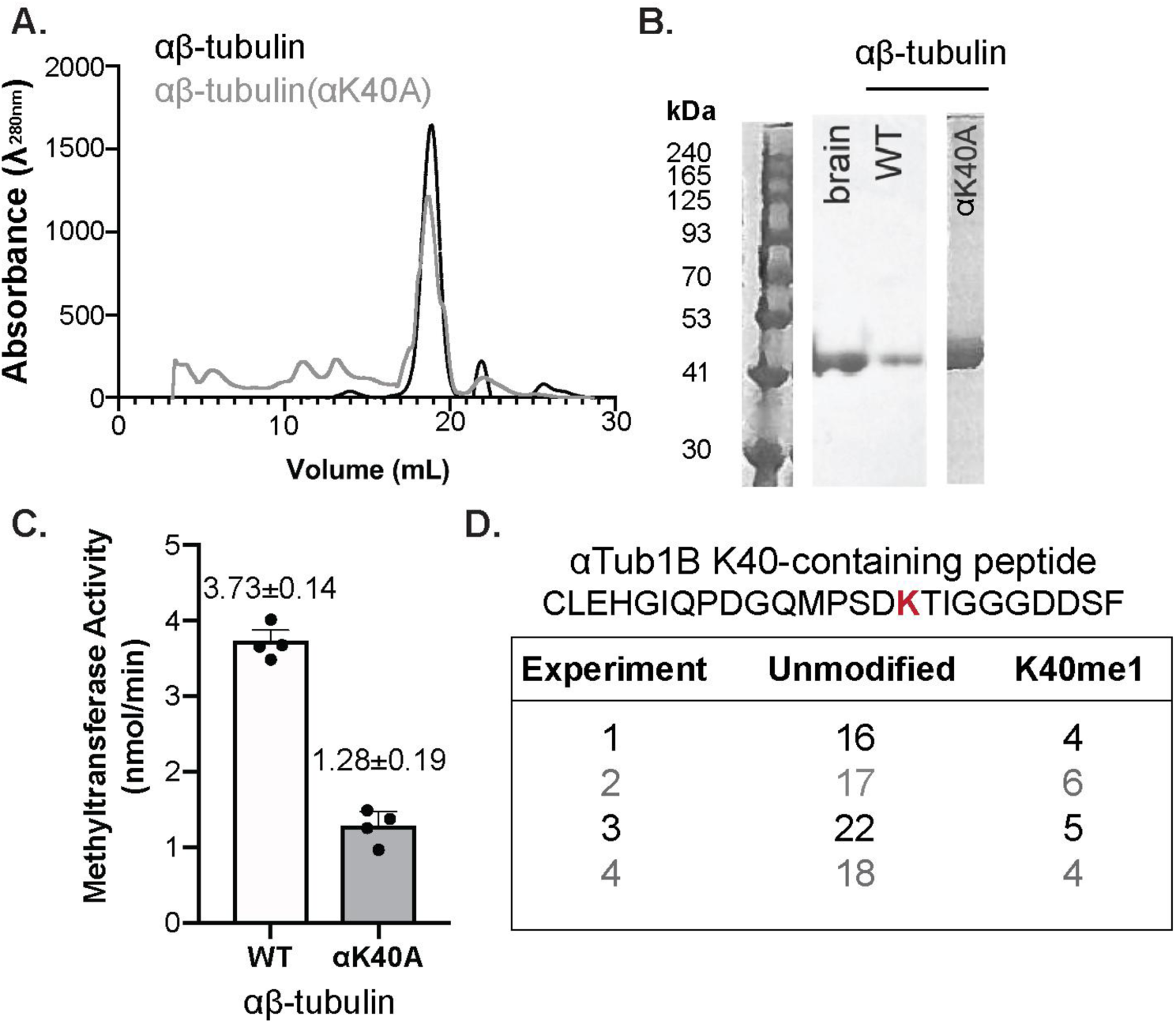
αTubK40 is the primary site for methylation by tSETD2. A,B) Recombinant single-isotype αTubA1B-βTub3B (αβ-tubulin) proteins, both WT and the αK40A mutant, were purified from Hi-5 insect cells and analyzed by A) size exclusion chromatography and B) Coomassie stained SDS-PAGE gel. C) Methyltransferase activity of tSETD2-FLAG incubated with 5 μM WT αβ-tubulin or mutant αβ-tubulin(αK40A) recombinant tubulins. Each dot indicates the average result from a single experiment and the bar represents the average±SD methyltransferase activity across n=4 experiments. D) Mass spectrometry analysis of tubulin peptides. Table shows the only peptide modified in the presence of both tSETD2-FLAG and SAM, followed by the number of peptides either unmodified or with K40 mono-methylated per experiment.

The ability of tSETD2-FLAG to methylate WT αβ-tubulin versus mutant αβ-tubulin(αK40A) dimers was then measured using the methyltransferase assay. We found that the tubulin methyltransferase activity of tSETD2-FLAG was decreased when provided with the mutant αβ-tubulin(αK40A) as a substrate (1.28±0.19 nmol/min) as compared to the methyltransferase activity towards WT αβ-tubulin (3.73±0.04 nmol/min) (Fig. 3C). This result suggests that although αTubK40 is the major site for methylation by tSETD2, there may be additional methylation sites on α- or β-tubulin. To identify potential methylation site(s), we carried out mass spectrometry analysis of methyltransferase reactions containing tSETD2-FLAG and WT or K40A mutant tubulins. As a negative control, experiments lacking the methyl donor SAM were carried out in parallel. For WT tubulin, we could only identify mono-methylation on αΜ0 (Figure 3D) and only in the presence of both tSETD2-FLAG and SAM. In the case of αK40A mutant tubulin, we were unable to identify any methylation sites on α- or β-tubulin (Fig 3D) even in the presence of both tSETD2-FLAG and SAM. These results suggest that αTubK40 is the only site methylated by tSETD2-FLAG.

### A R2510H SRI-domain mutation alters tSETD2’s ability to bind to and methylate tubulin

SETD2’s methylation of tubulin protein is dependent on its SET and SRI domains (19, 20). Clear cell renal cell carcinoma (ccRCC) mutations to the SET domain, R1625C, abolish SETD2 catalytic activity, but the molecular basis for reduced methylation of tubulin by R2510H in the SRI domain remains unclear. To investigate this, we generated and purified tSETD2(R2510H)-FLAG from HEK293 cells (Fig. 4A,B). As a control, we also generated and purified tSETD2-Flag containing a mutation in the catalytic SET domain, tSETD2(R1625C)-FLAG, (Fig 3 A,B) Using purified tSETD2 proteins and the methyltransferase assay, we confirmed that mutation of R1625C in the catalytic domain abolished the ability of tSETD2-FLAG to methylate both porcine brain tubulin protein and the control H3 peptide whereas mutation of R2510H in the SRI domain only abolished the ability of tSETD2-FLAG to methylate tubulin protein (Fig 4C).

**Figure 4:**
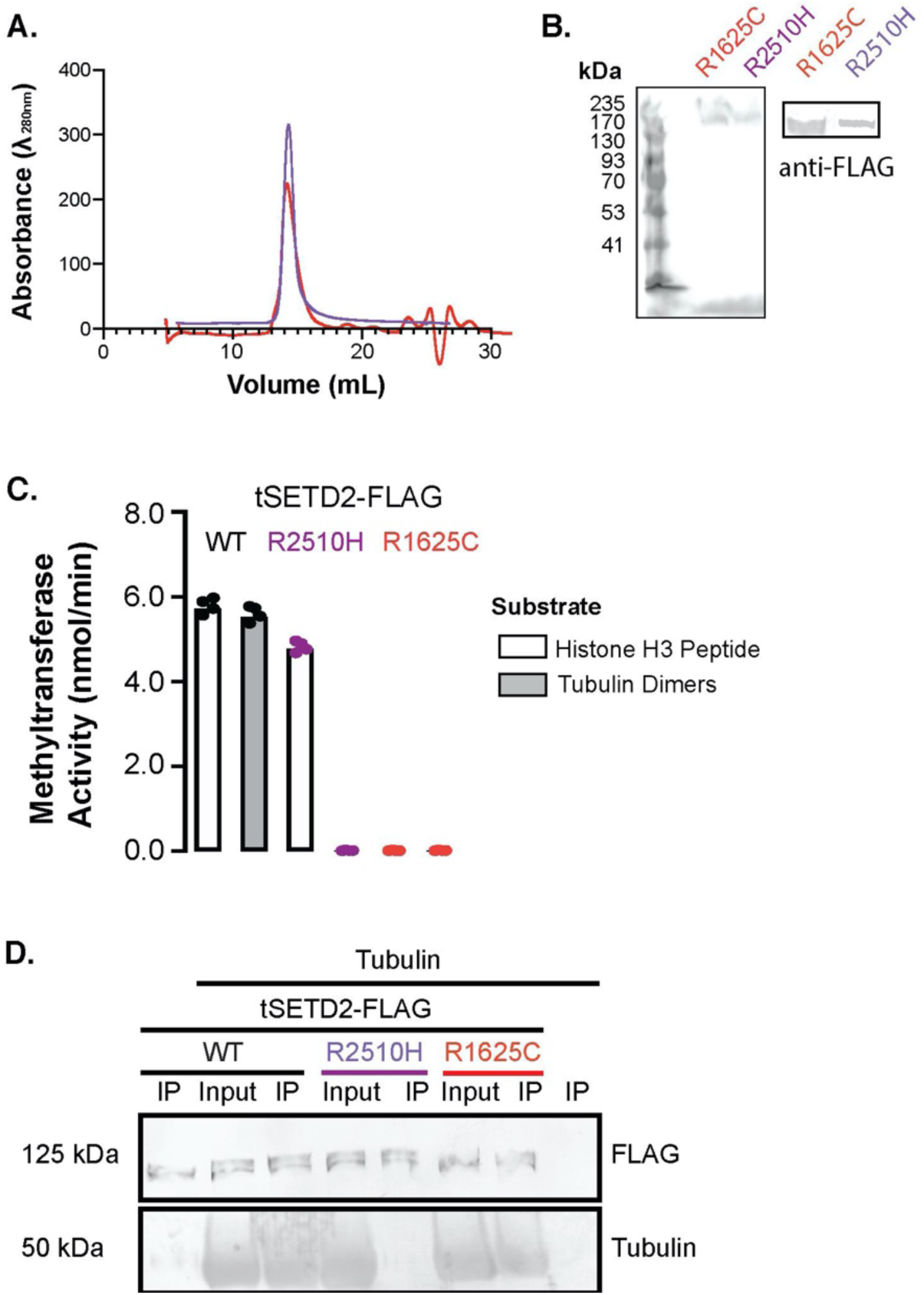
ccRCC-associated R2510H mutation in the SRI domain blocks binding and methylation of tubulin. A-B) tSETD2-FLAG protein with mutation of R1625C (red) or R2510H (purple) were purified from HEK293 cells and analyzed by A) size exclusion chromatography and B) Coomassie-stained gel (left) and anti-FLAG western blot (right). C) Methyltransferase activity of WT, R1625C, and R2510H tSETD2-FLAG against 5 μM histone-H3 peptide (white) or porcine brain tubulin (grey). Each dot indicates the average of a single experiment and the bar represents the average±SD methyltransferase activity across n=4 experiments. D) Co-immunoprecipitation of tubulin protein with WT, R1625C, or R2510H tSETD2-FLAG. The input and anti-FLAG pellet (IP) fraction were blotted with antibodies against the FLAG tag (top) and β-tubilin E7 (bottom).

We hypothesized that the SRI domain mutation reduced tubulin methylation by decreasing SETD2’s ability to bind to tubulin. To test this, we performed co-immunoprecipitation assays to assess the interaction of WT, R1625C, and R2510H tSETD2-FLAG with brain tubulin. We found that both WT tSETD2-FLAG and the SET domain mutant tSETD2(R1625C)-FLAG bound to tubulin, but that the tSETD2(R2510H)-FLAG SRI-domain mutation abolished the ability of tSTED2-FLAG to bind to tubulin (Fig. 4D). These results suggest that 1) the ccRCC-associated mutation of R2510H in the SRI domain abolishes the ability of tSETD2 to bind to and methylate tubulin as a substrate and 2) the SRI domain is critical for interaction with tubulin as a substrate.

### Distinct residues in the SRI domain allow substrate selection for tubulin or RNA Polymerase II

The SRI-domain of SETD2 has previously been shown to engage in protein-protein interactions with the highly phosphorylated C-terminal domain (CTD) repeat of RNA polymerase II (Pol II) for recruitment of SETD2 to chromatin during transcription (28, 34). Thus, our finding that mutation of R2510H in the SRI abolishes tubulin binding (Fig 4D) suggests that the SRI domain plays a major role in substrate recognition for both tubulin and RNA Pol II substrates. To test this, we took advantage of NMR-based studies that identified SRI residues impacted by binding to peptides mimicking the phosphorylated C-terminus of RNA Pol II (28, 34). We targeted these residues and generated tSETD2-FLAG variants with alanine mutations to V2483, F2505, K2506, R2510, and H2514 (Fig 5A). We also generated a construct lacking the SRI-domain SRI) as a control (Fig 5A). The mutant and deletion variants of tSETD2-FLAG were purified from mammalian cells (Fig. 5B,C).

**Figure 5:**
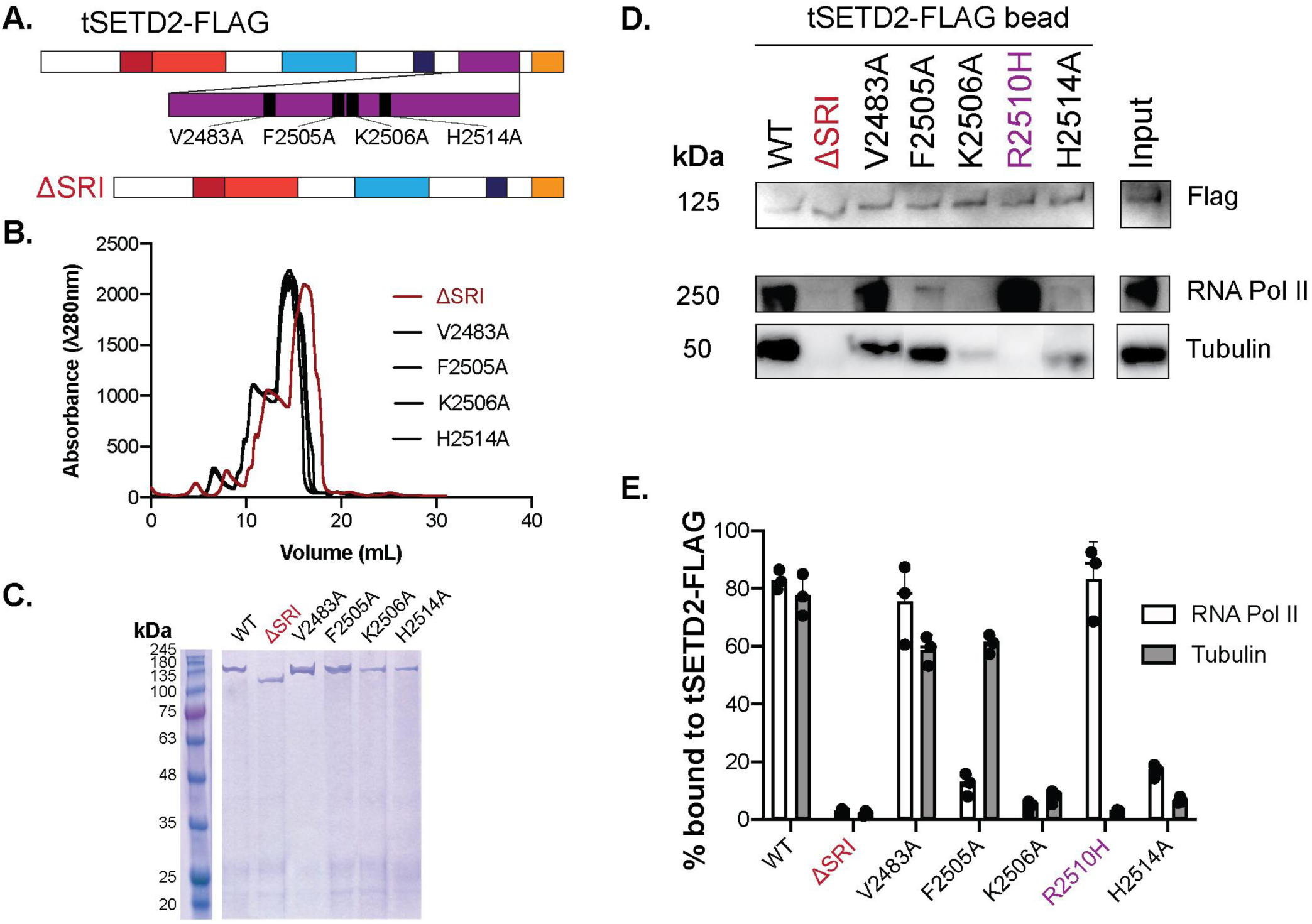
Residues in the SRI-domain of SETD2 distinguish tubulin and RNA-Pol II binding. A) Schematic of point mutations or deletion of the SRI domain in tSETD2-FLAG. B-C) Purification of tSETD2-FLAG containing mutations in or deletion of the SRI-domain. The proteins were analyzed by B) size-exclusion chromatography and C) Coomassie-stained gel. D,E) Purified WT, ΔSRI, or mutant versions of tSETD2-FLAG bound to anti-Flag beads were incubated with either tubulin protein or HEK293 cell lysate. The presence of tSETD2-FLAG variant, tubulin, and Pol II in the bead pellet was analyzed by western blotting with antibodies to the FLAG tag, α-tubulin, and Pol II, respectively. D) Representative image. The far-right column shows the input for the reaction with WT tSETD2-FLAG. E) Quantification of RNA Pol II (white) and porcine brain tubulin (grey) co-pelleting with tSETD2-FLAG as a percentage of the input reaction. Each dot indicates the percent bound from one experiment and the bar represents the average±SD across n=3 experiments.

We tested the ability of the purified mutant and deletion tSETD2-FLAG proteins to bind to RNA Pol II and tubulin using the co-immunoprecipitation assay. The tSETD2-FLAG proteins bound to Anti­FLAG beads were incubated with either porcine brain tubulin or HEK293 cell lysates containing endogenous RNA Pol II. As expected, deletion of the SRI domain SRI) abolished the ability of tSETD2-FLAG to bind to both RNA Pol II and tubulin substrates (Fig. 5D,E). Interestingly, the SRI-domain mutants varied in their ability to bind to the two substrates. The V2483A mutant retained binding to both substrates, the F2505A retained tubulin but not RNA Pol II binding, the R2510H mutant retained RNA Pol II but not tubulin binding, and the K2506A and H2514A mutants lost the ability to bind to both tubulin and RNA Pol II (Fig. 5D,E). These results indicate that the SRI domain is involved in substrate recognition and that distinct residues in the SRI domain contribute to recognition of different substrates.

### SETD2 recognizes tubulin via α-tubulin’s C-terminal tail (CTT)

The interaction between SETD2 and RNA Pol II involves the negatively-charged phosphorylated C-terminal repeat domains of Pol II (28, 34)). This led us to hypothesize that SETD2 recognition of tubulin involves similar charge-charge interactions, particularly between positively-charged residues in the SETD2 SRI domain (Fig 5D,E) and the negatively-charged C-terminal tails (CTTs) of α- and/or β-tubulin. To test this, we used our recombinant single-isotype tubulin to generate αβ-tubulin proteins containing truncations of the CTTs of either α-tubulin [αβ-tubulin (ΔαCTT)] or β-tubulin [αβ-tubulin(ΔβCTT)] and purified the tubulin dimers from insect cells (Fig. 6A,B).

**Figure 6:**
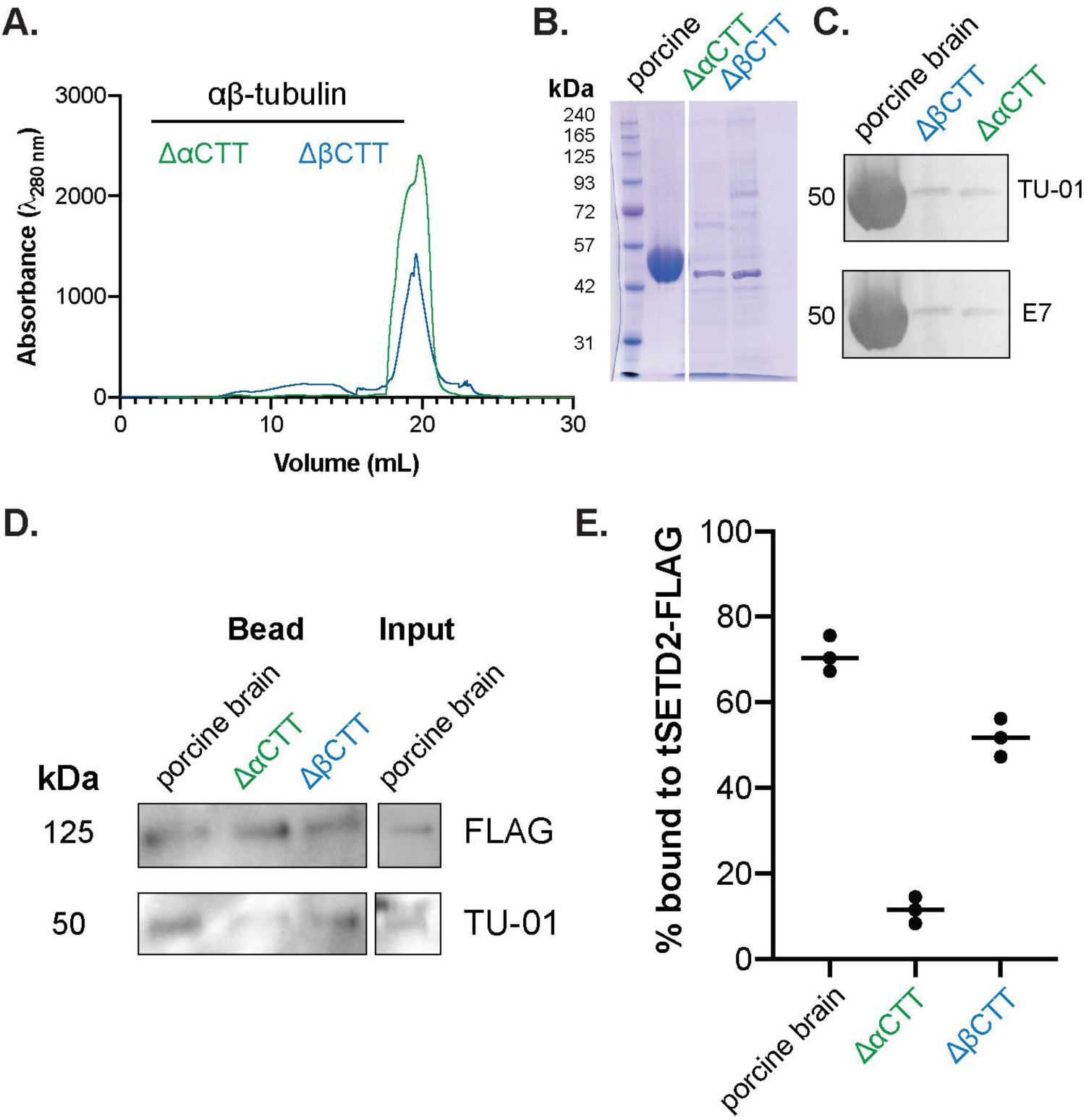
tSETD2-FLAG binds to the C-terminal tail (CTT) of α-tubulin. A,B) Tailless forms of recombinant tubulin, αβ-tubulin(ΔαCTT) and αβ-tubulin(ΔβCTT), were purified from Hi-5 insect cells and analyzed by A) size exclusion chromatography and B) Coomassie-stained gel. C) Western blot of porcine brain tubulin, purified αβ-tubulin(ΔαCTT), and purified αβ-tubulin(ΔβCTT) with antibodies that recognize the N-terminus of α-tubulin (TU-01, top) or the CTT of β-tubulin (E7, bottom). D) Co-immunoprecipitation of tSETD2-FLAG with porcine brain tubulin, αβ-tubulin(ΔαCTT), or αβ-tubulin(ΔβCTT). Shown are western blots of the bead pellets with antibodies to tSETD2-FLAG (FLAG, top) and the N-terminus of α-tubulin (TU-01, bottom). The far-right column shows the input for the reaction with porcine tubulin. E) Quantification of amount of tubulin co-pelleting with tSETD2-FLAG compared to input. Each dot represents the percent bound from one experiment and the line represents the average across three independent experiments.

We tested the ability of tSETD2-FLAG to bind to the tailless tubulins using a co-immunoprecipitation assay. As most anti-tubulin antibodies recognize the CTTs, we first identified conditions for detecting the tailless tubulins by western blotting. For this, we used a polyclonal antibody TU-01 whose epitope resides in the N-terminus of α-tubulin. As expected, the TU-01 antibody was able to detect both αβ-tubulin(ΔαCTT) and αβ-tubulin(ΔβCTT) proteins by western blotting whereas an antibody against the CTT of β-tubulin (E7) failed to recognize the αβ-tubulin(ΔβCTT) protein (Fig 6C). Immobilized tSETD2-FLAG was incubated with either αβ-tubulin(ΔαCTT) or αβ-tubulin(ΔβCTT) tubulin proteins. Whereas t-SETD2-FLAG pulled down porcine brain and αβ-tubulin(ΔβCTT) tubulins, it was unable to co-precipitate αβ-tubulin (ΔαCTT) tubulin (Fig 6D,E). From these experiments, we conclude that tSETD2-FLAG binds to tubulin via the negatively-charged CTT of α-tubulin.

## Discussion

In this study, we utilized *in vitro* biochemical reconstitution with recombinant proteins to determine how SETD2 recognizes and methylates tubulin. By exploiting known tubulin-targeting agents, we found that SETD2 preferentially methylates the dimeric form of tubulin vs. microtubule polymers. Interestingly, our work indicates that SETD2’s SRI domain makes electrostatic interactions with the CTT of α-tubulin in a mechanism distinct from RNA Pol II targeting, suggesting that this interaction positions the SET domain for methylation of residue K40 of α-tubulin (Fig. 7).

**Figure 7:**
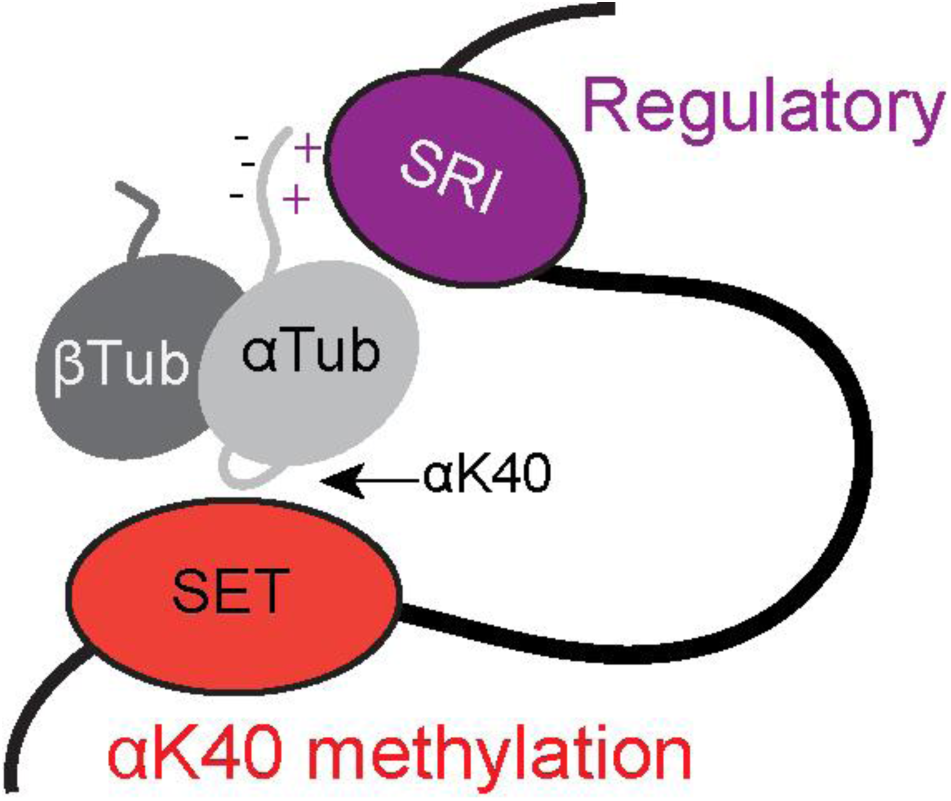
Proposed model of tubulin methylation by tSETD2. The SRI-domain (purple) recognizes the negatively-charged CTT of α-tubulin (grey) and positions the SET domain (red) for methylation of α-tubulin at the K40.

### SETD2 methylation is restricted to K40 of α-tubulin

Our mass spectrometry analysis identified α-tubulin K40 as the only detectable site of methylation by tSETD2 on the recombinant single-isotype αβ-tubulin. This confirms and extends the previous work where tri-methylation was detected at this site in mammalian cells (19). In our mass spectrometry experiments, we were only able to detect a mono-methylation mark on αΚ40 whereas in cells, trimethylated tubulin was detected (19). The canonical SETD2 activity is with di-methylated histone substrates. While this suggests that mono- or di-methylated tubulin could be a preferred substrate for SETD2, it is known that SETD2 can mono-methylate substrates (35), but typically will di-or tri-methylate substrates. As such, tubulin methylation may require priming by other methyltransferases before SETD2 makes its mark in cells.

Although mutation of K40A did not completely abolish methyltransferase activity measured in the fluorescence-based assay, we were unable to detect any other methylated peptides for α- or b-tubulin by mass spectrometry. It seems unlikely that the residual methyltransferase activity comes from tSETD2 itself as the auto-methylation activity is subtracted from our measurements. It is possible that SETD2 is able to methylate another tubulin residue that we have been unable to detect in our mass spectrometry analysis, perhaps because the methylated peptide does not ionize well or the amount is below the limit of detection. It is also possible that tSETD2-FLAG or a co-purifying methyltransferase enzyme from HEK293 cells is able to methylate a contaminating protein that co-purifies with tubulin from the insect cells and/or with tSETD2-FLAG from the HEK293 cells. Further work will be required to discern between these possibilities.

### Crosstalk between acetylation and methylation of α-tubulin K40

The K40 residue resides on a flexible loop of α-tubulin that is located within the lumen of a polymerized microtubule (36, 37), and could be accessible to modifying enzymes when tubulin is in either the soluble dimer or microtubule form. However, given the large size of the SETD2 protein (∼290 kDa) and the restricted size of the microtubule lumen (∼17 nm diameter), it has been puzzling how SETD2 could access the K40 residue within the microtubule lumen (19). We demonstrate that tSETD2 has higher methylation activity towards soluble tubulin over microtubules (Fig. 2). Binding of SETD2 to soluble tubulin requires the SRI domain of tubulin and the CTT tail of α-tubulin, an interaction that likely positions the SET domain for methylation of K40.

The K40 residue of α-tubulin is also known to be acetylated by α-tubulin acetyltransferase (αTAT) (38, 39). Whether SETD2 and αTAT compete for access to the K40 residue has been unclear. Given our results demonstrating a higher activity of SETD2 towards soluble tubulin and recent work demonstrating that αTAT1 preferentially acetylates polymerized microtubules and enters the microtubule lumen (39–43), the two enzymes appear to work on tubulin in different contexts. Specifically, SETD2 preferentially methylates soluble tubulin whereas αTAT preferentially acetylates polymerized tubulin. The interplay between these enzymes is likely to play a role in specific cellular events. For example, elevated levels of α-tubulin acetylation correlate positively with metastatic potential (44) and αTAT inhibits cancer cell motility (45), whereas SETD2-mediated tubulin methylation corresponds with genomic stability and correct mitotic spindle formation (19, 20).

Interestingly, both SETD2 (this work) and αTAT (39) display slow enzymatic rates toward the tubulin substrate *in vitro.* SETD2 also shows similar low k_cat_ values towards purified nucleosome substrate (31). In a histone methylating context, SETD2 is typically recruited by other protein complexes (e.g. IWS1, SPT6, and RNA Pol II) (46, 47) suggesting that substrate binding and enzyme kinetics could be higher in cells.

### The role of the SETD2 SRI domain in substrate binding

Previous work demonstrated that the ccRCC-associated mutation R2510H in the SRI domain abolished the ability of SETD2 to methylate tubulin but not histone H3 (18, 19). We provide a mechanism for this finding by demonstrating that the R2510H mutation abolishes the ability of tSETD2 to bind to tubulin (Figs. 4,5) but has no effect on its ability to bind to interact with RNA Pol II from cell lysate (Fig 5). That the SRI domain is required for binding to tubulin appears to contradict previous work suggesting that the SET domain is sufficient for tubulin binding (19). These discrepancies could be due to the use of different constructs and expression systems, as well as differences in experimental protocols. The fact that the interaction of RNA Pol II is less sensitive to the R2510H mutation than tubulin may be related to the sequence and charge of these binding segments. Specifically, the CTD of Pol II contains 22 heptapeptide (YSPTSS) repeats and is highly phosphorylated whereas the CTT of α-tubulin contains a short sequence of negatively-charged residues. Thus, the interaction between the SRI domain of SETD2 with RNA Pol II may better tolerate the R2510H mutation than the interaction of the SRI domain with α-tubulin. If so, caution should be taken when studying interactions with RNA Pol II CTD peptides containing fewer repeats which could produce different results than using full-length RNA Pol II. Nevertheless, these results provide strong support for the hypothesis that genomic instability in ccRCC can be driven by tubulin-dependent functions of SETD2 contributed by the SRI domain.

By mutagenesis of the SRI domain based on structural information in the literature (28, 48), we were able to identify residues that further distinguish the ability of SETD2 to bind to tubulin versus Pol II. We demonstrate that the ability of tSETD2 to bind to tubulin requires not only the SRI domain residue R2510, but also K2506 (Fig 5). In contrast, the ability of tSETD2 to bind Pol II is influenced by F2505, K2506, and H2514. These results are largely consistent with previous studies using NMR structures of the SRI domain and titration experiments with phosphorylated Pol II peptides that implicated residues V2483, F2505, K2506, R2510, H2514 in forming part of the SRI-Pol II binding interface (28, 34, 49). However, while the isolated SRI domain with the R2510H point mutant showed decreased binding to RNA Pol II CTD peptides (28), our results suggest that R2510H does not decrease the ability of tSETD2-FLAG to pull-down Pol II from cell lysates (Fig. 5C). These differences are likely due to differences in experimental conditions including the use of an isolated SRI domain versus tSETD2 and the use of Pol II peptides versus full-length CTD as mentioned above.

### Recognition of the α-tubulin CTT by SETD2

Our results also demonstrate that tSETD2 requires the CTT of α-tubulin, a negatively-charged region of the protein, for binding (Fig 6). This is interesting given previous work demonstrating that the SRI domain of SETD2 interacts with the phosphorylated, and thus negatively-charged, C-terminal repeat domains of Pol II (34, 48, 50). It is thus tempting to speculate that charge-charge interactions are key for recognition of substrates by the SRI domain of SETD2. However, actin has recently been shown to be methylated by SETD2 (51) and unlike tubulin and RNA Pol II, actin does not have a long flexible negatively-charged tail or loop within its structure. However, in these studies, actin methylation by SETD2 required other binding partners, namely the Huntingtin protein, HTT, and the actin binding adaptor protein HIP1R. Future work will be required to delineate the mechanism by which SETD2 recognizes actin and other substrates.

While our results suggest that SETD2 does not compete with α TAT for modification of the K40 residue, it likely competes with other tubulin-interacting proteins and/or modifying enzymes that target the CTT. The flexible CTTs of tubulin subunits extend from the surface of the microtubule and form a negatively-charged surface that appears to serve as a recognition site for a large number of microtubule associated proteins. For example, tubulin tyrosine ligase (TTL) makes critical electrostatic interactions with α-tubulin residues E445, E446, and E447 in order to align the CTT within the active site (52). The VASH-SVBP complex that removes the terminal tyrosine from α-tubulin, clear density of the CTT remains unresolved by both x-ray crystallography and cryo electron microscopy (53, 54) makes electrostatic interactions with a compound, epoY, that mimics the tubulin CTT residues (55). Similarly, the tubulin CTTs are essential for microtubule recognition by tubulin tyrosine ligase-like 7 (TTLL), an enzyme that adds glutamate chains to the CTTs of both α- and β-tubulin (56, 57). As such, our work adds SETD2 to the growing list of tubulin modifying enzymes that recognizes tubulin CTTs.

### Implications in cancer pathologies

Our findings provide further support for a role of SETD2 in writing both the histone and tubulin codes. As mutations in SETD2 continue to be identified in a growing list of tumor types (20, 58), it will be important to discern the relative roles of histone versus tubulin methylation in contributing to the underlying mechanisms of particular cancer phenotypes. A lack of tubulin methylation has been documented to result in multipolar spindles, genomic instability, and micronuclei formation (18–20). A better understanding between histone and tubulin methylation by SETD2 could also drive anti-cancer drug development and thus could provide new therapeutic targets to help cancer patients. Interestingly, the histone H4 lysine 20 methylating enzyme SET8, along with transcription factor LSF, has been identified as a modifier of α-tubulin at K311 although the cellular implication of loss of K311 methylation remains to be determined (59). Other tubulin PTMs have been shown to impact mitotic progression and may also underlie cancer phenotypes. Recent work has shown that a disruption of α-tubulin detyrosination leads to reduced chromosome congression and increased errors of kinetochore-microtubule attachment (60-63). Further investigations into tubulin PTMs and their modifying enzymes are required to understand the nuanced interaction between histone and tubulin modifications in cells and the implications for cancer progression.

## Acknowledgements

We would like to thank Venkatesha Basrur, Alexey Nesvizhskii, and the Proteomics Resource Facility (PRF) at the University of Michigan Department of Pathology for conducting and analyzing the mass spectrometry experiments. We also acknowledge the generous gifts of αTub1B/βTub3 plasmids and advice on protein expression and purification from Tarun Kapoor and members of his laboratory at Rockefeller University. We also thank Dan Sackett at the National Institutes of Health for advice on podophyllotoxin. S.K. is funded by Graduate Assistance in Areas of National Need (P200A150164) and the Chemical Biology Interface training grant from the NIH (5T32GM008597-22). This work was supported by NIH grant UL1TR002240 to the Michigan Institute for Clinical and Health Research (MICHR) and by NIH grants R01GM070862 and R35GM131744 to KJV. The authors do not have any conflicts of interest.

## Materials and Methods

### Plasmids

An active truncated SETD2 construct (1418–2564) with a FLAG affinity tag (tSETD2-FLAG) in the pInducer vector for mammalian expression was generated by the Walker Lab. Single isoform αTub1B/βTub3 plasmid cDNA encoding Homo sapiens α-tubulin 1B (NP_006073.2), β-tubulin 3 (NM_178012.4) in pFastBac Dual vector (ThermoFisher 10712024) was obtained from the Kapoor lab for insect cell expression (23). Point mutations and domain deletions were generated using QuickChange site-directed mutagenesis with Q5 Polymerase (NEB). All plasmids were verified by DNA sequencing.

### Purification

#### SETD2

tSETD2-FLAG was transfected into HEK 293 Freestyle cells with FectoPRO transfection reagent (Polyplus, 10118-444) and cells were harvested 48 hours later at 5K rpm for 15 mins (Beckman JLA 8.1).The pellet was suspended in lysis buffer (50 mM HEPES pH 7.5, 50 mM MgCl2, 150 mM NaCl, cOmplete protease inhibitor tablet (Sigma Aldrich, 4693159001) and cells were lysed with 20 strokes of a dounce homogenizer. This was ultracentrifuged (Beckman Ti70 337922) at 40K rpm for 40 mins and the supernatant was filtered with 1.0 um glass fiber filter (Pall Laboratory) and incubated with FLAG M2 affinity beads (Sigma Aldrich) equilibrated in lysis buffer for 3 hours. Beads were rinsed with 3 column volumes of wash buffer (50 mM NaPi pH 7.2, 150 mM NaCl, 5 mM BME), 3 CV salt buffer (wash buffer at 500 mM NaCl), and again with wash buffer before elution buffer (wash buffer with 300 ng 3x-FLAG peptide (Sigma Aldrich)) was added and incubated with beads overnight. Eluent was then run over ion exchange column (DEAE Sepharose, GE Life Sciences) on a 0-75% salt buffer gradient, and size exclusion chromatography (Superose 6 Increase 10/300, Fisher Scientific) with gel filtration buffer (50 mM sodium phosphate (NaPi) pH 7.2, 150 mM NaCl, 5 mM BME, 5% glycerol). Fractions were pooled and concentrated down with an Amicon Ultra 100K MWCO centrifugal filter unit and snap frozen in liquid nitrogen and stored at −80C.

#### aTub1BTub3 tubulin

Purification was as previously described (23, 64). Briefly, the Bac-to-Bac system (Life Technologies) was used to generate recombinant baculovirus in SF9 cells. High Five cells (Thermo Fisher, B85502), grown to 3 million cells/ml in Lonza Insect XPRESS (Fisher Scientific, BW12-730Q), were infected with P3 viral stocks at ∼10 mL/L. Cells were cultured in suspension at 27°C and harvested at 60 hours after infection. The following steps were done on ice or at 4°C. Cells were lysed in an equal volume of lysis buffer (50 mM HEPES, 20 mM imidazole, 100 mM KCl, 1 mM MgCl2, 0.5 mM β-mercaptoethanol, 0.1 mM GTP, 3 U/ml benzonase, 1X Roche Complete EDTA-free protease inhibitor, pH 7.2) by dounce homogenization (20 strokes) and the homogenate was centrifuged at 55,000 rpm in a Type 70 Ti rotor (Beckman Coulter) for 1 hr. The supernatant was then filtered through a 0.22 μm membrane (Fisher Scientific, 09740113) and loaded onto a 5 ml HisTrap HP column (GE life science 17-5247-01) pre-equilibrated with lysis buffer. The column was washed with 35 ml lysis buffer until the UV absorption reached baseline, then eluted with nickel elution buffer (1X BRB80 (80 mM PIPES, 1mM MgCl2, 1mM EGTA), 500 mM imidazole, 0.2 mM GTP, 2 mM β-mercaptoethanol, pH 7.2). The fractions containing proteins were pooled, diluted 3-fold with lysis buffer and loaded one 5 ml StrepTrap HP column (GE life science 29-0486-53). The column was washed with 25 ml 66% lysis buffer + 33% nickel elution buffer, 25 ml of wash buffer 1 (1X BRB80, 1 mM β-mercaptoethanol, 0.1 mM GTP, 0.1 % Tween-20, 10% glycerol, pH 7.2), and 25 ml of wash buffer 2 (1X BRB80 1 mM β-mercaptoethanol, 0.1 mM GTP, 10 mM MgCl2, 5 mM ATP, pH 7.2). The bound protein was then eluted with ∼5 ml StrepTrap elution buffer (1XBRB80, 20 mM Imidazole, 2 mM β-mercaptoethanol, 0.2 mM GTP, 3 mM desthiobiotin, pH 7.2). The StrepTrap eluate was mixed with 4 mg of previously purified TEV protease (∼8 mg/ml stored in 40 mM HEPES, 150 mM KCl, 30%(w/v) glycerol, 1 mM MgCl2, 3 mM β-mercaptoethanol, pH 7.5) and incubated for 2 hr on ice. The TEV-digested protein solution was concentrated with an Amicon Ultra 50K MWCO centrifugal filter unit (Millipore UFC901024) to 2 mL, and loaded onto a Superdex 200 Increase 10/300 GL column equilibrated in size-exclusion buffer (1XBRB80, 5%(w/v) glycerol, 0.2 mM GTP, 2 mM β-mercaptoethanol, pH 6.8). Tubulin eluted at ∼15 ml and was concentrated to >3 mg/ml with an Amicon Ultra 50K MWCO centrifugal filter unit. The purified tubulin was snap frozen in liquid nitrogen and stored at −80°C. Tail-less tubulin eluted at ∼18 mL and concentrated with an Amicon Ultra 30K MWCO centrifugal filter unit, but otherwise was purified the same way.

### Methyltransferase assay

The activity of tSETD2-FLAG constructs was measured using a Methyltransferase Fluorescence Assay Kit (Cayman Chemical, 700150). This continuous enzyme-coupled assay continuously monitors SAM-dependent methyltransferases by generating a fluorescent compound, resorufin, from the reaction product, AdoHcy. Fluorescence is analyzed with an excitation wavelength of 530-540 nm and an emission wavelength of 585-595 nm using a PHERAstar Plate Reader (BMG Labs). A standard curve of resorufin concentration and fluorescence was used to determine concentration-dependent fluorescence. The initial velocities of the reaction curves were obtained by linear-regression, and then were plotted in a concentration-dependent manner to obtain Michaelis-Menten plots, and as such values for Km and vmax (Prism Version 8.1.1).

#### Dimer or microtubule stabilization

Microtubule polymerization was inhibited by the addition of 50 μM of podophyllotoxin provided by Dan Sackett (Millipore Sigma, P4405). Microtubule stabilization occurred with 100 μM taxol (Cytoskeleton Inc, TXD01). Stocks were made at 2 mg/mL and then diluted to perform methyltransferase assay in BRB80.

### Mass Spectrometry

Purified single isoform tubulin and tSETD2-FLAG were incubated at molar ratio of 5:1 with excess S-adenosylmethionine (Sigma Aldrich, 86867-01-8) for 2 hours at room temperature. The mixture was then digested with fresh Chymotrypsin (Sigma Aldrich, 11418467001) and put onto a Thermo Scientific mass spectrometer with an Orbitrap Fusion Tribrid with ETD and a Q Exactive HF equipped with a nano-LC system (Dionex RSLC-nano). Analysis of PTMs was conducted with PEAKS X software.

### Pull-down assay

FLAG M2 beads were blocked with 3% BSA in PBS for 1 hour and equilibrated in the reaction buffer (tSETD2-FLAG size-exclusion buffer, described previously). tSETD2-FLAG protein (WT or variants) was added at 20 μΙ with a putative binding partner for 2 hours in the presence of SAM. For tubulin, 0.25 mg/mL of porcine tubulin (Cytoskeleton, Inc.) was used, and for RNA Polymerase II, 10 μL of HEK293 Freestyle clarified lysate was used. Beads were spun down and the supernatant was collected as the fraction of unbound substrate. The beads were then resuspended in the reaction buffer to the total reaction volume, and the same amount of supernatant and beads were added to SDS-PAGE gel. Analysis of binding was conducted by western blot with the following antibodies: anti-FLAG (Sigma Aldrich, A9469, 1:1000), anti-tubulin antibodies E7 (DSHB, AB_528499, 1:1000) and/or TU-01 (BioLegend **625902**, 1:1000), and anti-RNA Polymerase II (Abcam, ab193468, 1:1000), with secondary antibody anti-mouse (Enzo Life Science, ADI-SAB-100-J, 1:1000) or anti-rabbit (Enzo Life Science, ADI-SAB-300-J, 1:1000), respectively. Binding was quantified by measuring the background-subtracted intensity of each band with Fiji ImageJ (65) as a fraction of the input intensity. Each experiment was performed three times, independently.

### Immunohistochemistry

COS7 (ATCC, CRL-1651) cells transiently expressing tSETD2-FLAG constructs (Lipofectamine 2000 and OptiMEM) were fixed with 4% formaldehyde in PBS, treated with 50 mM NH4Cl in PBS to quench unreacted formaldehyde and permeabilized with 0.2% Triton X-100 in PBS. Subsequently, cells were blocked in blocking solution (0.2% fish skin gelatin in PBS). Primary antibodies tubulin (DSHB, AB_528499, 1:2000) and FLAG (Abcam, ab205606, 1:2000), and secondary antibodies were applied in blocking solution at room temperature for 1 h each, washing in between with blocking solution. Nuclei were stained with 10.9 μΜ 4’,6-diamidino-2-phenylindole (DAPI) and cover glasses were mounted in ProlongGold (Life Technologies). Cells were incubated 3x for 5 min in blocking solution to remove unbound antibodies. Images were collected on an inverted epifluorescence microscope (Nikon TE2000E) equipped with a 60x, 1.40 numerical aperture oil-immersion objective and a 1.5x tube lens on a Photometrics CoolSnapHQ camera driven by NIS-Elements (Nikon) software.

### Microtubule polymerization

#### GMPCPP-stabilized

We first prepared α/β3 microtubule seeds. Tubulin was thawed, mixed with GMPCPP (final 1.5 mM), diluted to ∼ 1.5 mg/ml with 1XBRB80+5% glycerol, centrifuged at 90,000 rpm for 10 min at 4°C (TLA120.1 Beckman Coulter), and then polymerized by incubation at 37°C for 30 mins. The microtubules were pelleted at 90,000 rpm for 10 min at 37°C (TLA120.1 Beckman Coulter) and re-suspended in warm (37°C) 1XBRB80 supplemented with 1 mM TCEP. Next, we used these microtubule seeds to polymerize GTP-bound α-π. Another aliquot of recombinant tubulin was thawed, diluted to a final concentration ∼3 mg/ml (1XBRB80, 33% glycerol, 1 mM GTP), and spun at 90,000 rpm for 10 min at 4°C (TLA120.1 Beckman Coulter). After incubation at 37°C for 2 mins, the supernatant was mixed with GMPCPP-seeds from the prior step and then incubated at 37°C for 30 mins followed by centrifugation at 90,000 rpm for 10 min at 37°C (TLA120.1 Beckman Coulter). The microtubule pellets were rinsed twice with 100 μ! warm (37°C) EM buffer (1X BRB80, 1 mM DTT, 0.1 mM ATP, 0.05% Nonidet P-40) before suspending in 30 μl cold EM buffer and then incubated on ice for 1 hr. After a centrifugation at 90,000 rpm for 10 min at 4°C, the supernatant containing depolymerized GDP-tubulin (∼2 mg/ml, measured by Bradford assay) was mixed with GMPCPP (final 2 mM) and then incubated on ice for 10 mins. After an incubation at 37°C for 2 mins, the protein solution was mixed with 30 μl warm (37°C) EM buffer followed by 37°C incubation for another 1 hr. The polymerized GMPCPP-microtubules were pelleted by 90,000 rpm for 10 min at 37°C (TLA120.1 Beckman Coulter) and suspended in warm (37°C) EM buffer.

## Supplemental Figures

**Figure S1:**
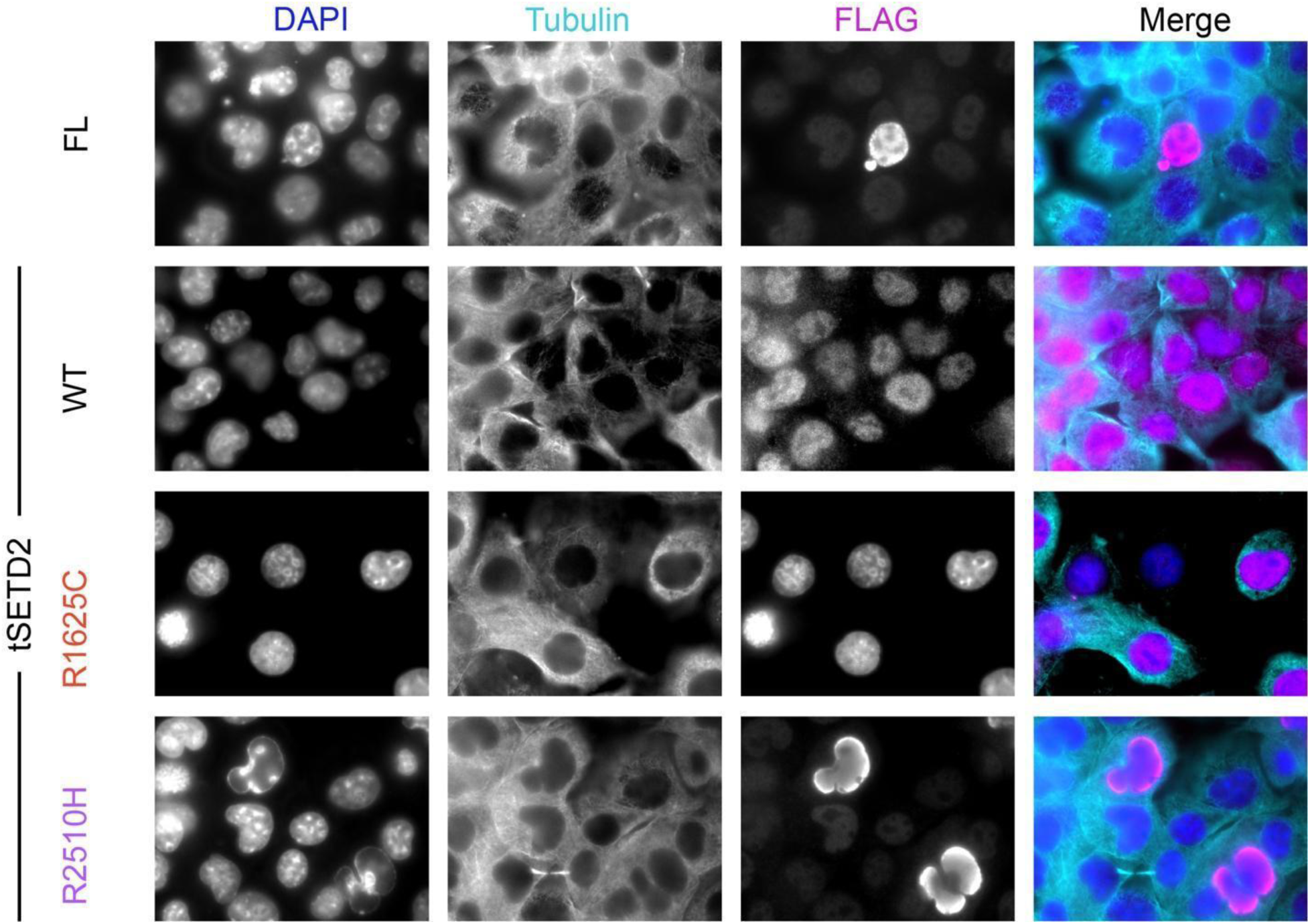
Localization of tSETD2-FLAG constructs in interphase cells. COS-7 cells transiently expressing (top row) FL SETD2-FLAG, (second row) tSETD2-FLAG, (third row) tSETD2(R1625C)-FLAG or (bottom row) tSETD2(R2510R)-FLAG were fixed and stained for DAPI (DNA, dark blue) and with antibodies against β-tubulin E7 (teal), and the FLAG tag (magenta).

**Figure S2:**
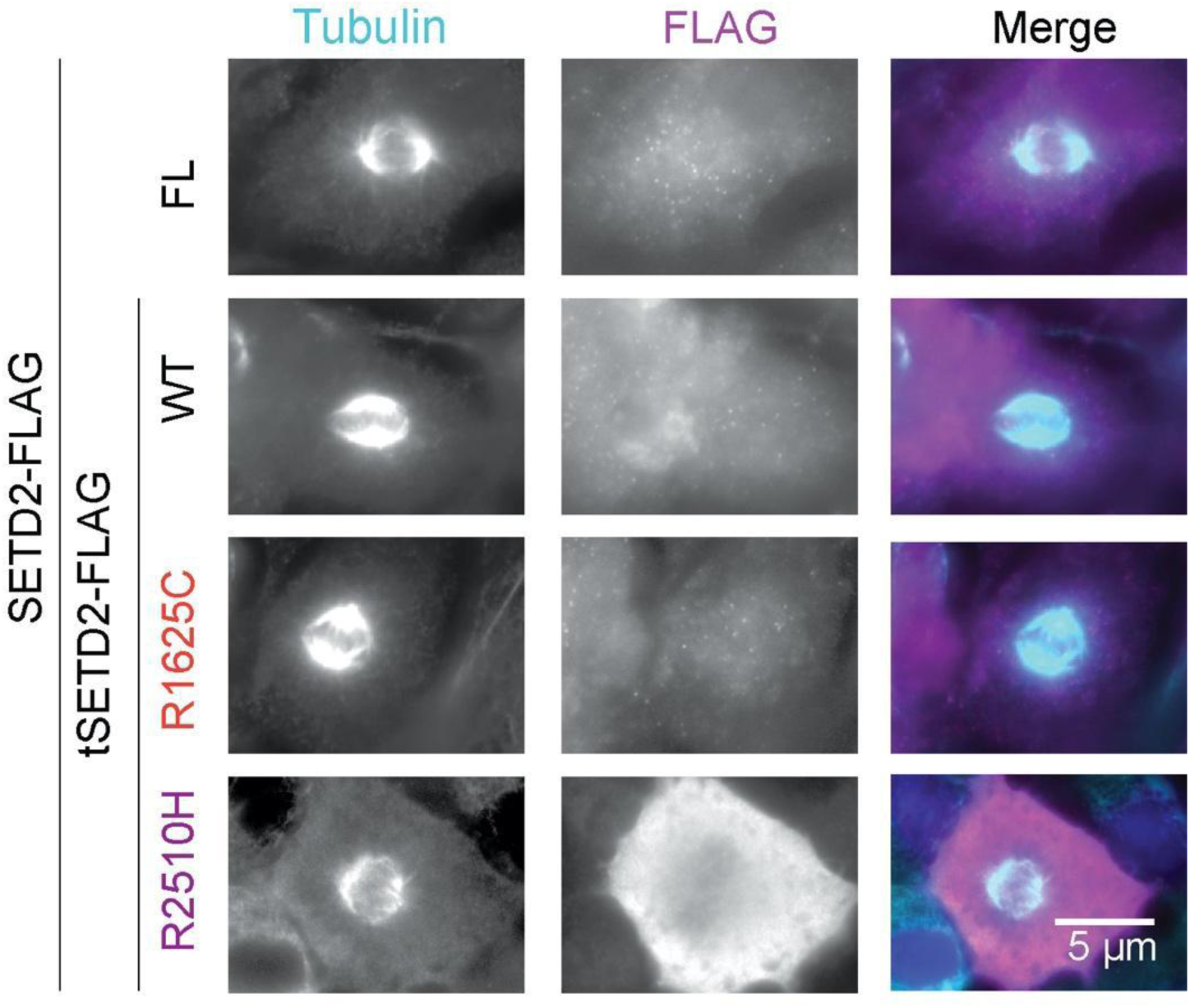
Localization of tSETD2-FLAG constructs in mitotic cells. COS-7 cells transiently expressing (top row) FL SETD2-FLAG, (second row) tSETD2-FLAG, (third row) tSETD2(R1625C)-FLAG or (bottom row) tSETD2(R2510R)-FLAG were fixed and stained with antibodies against α-tubulin (blue), and the FLAG tag (magenta).

**Figure S3:**
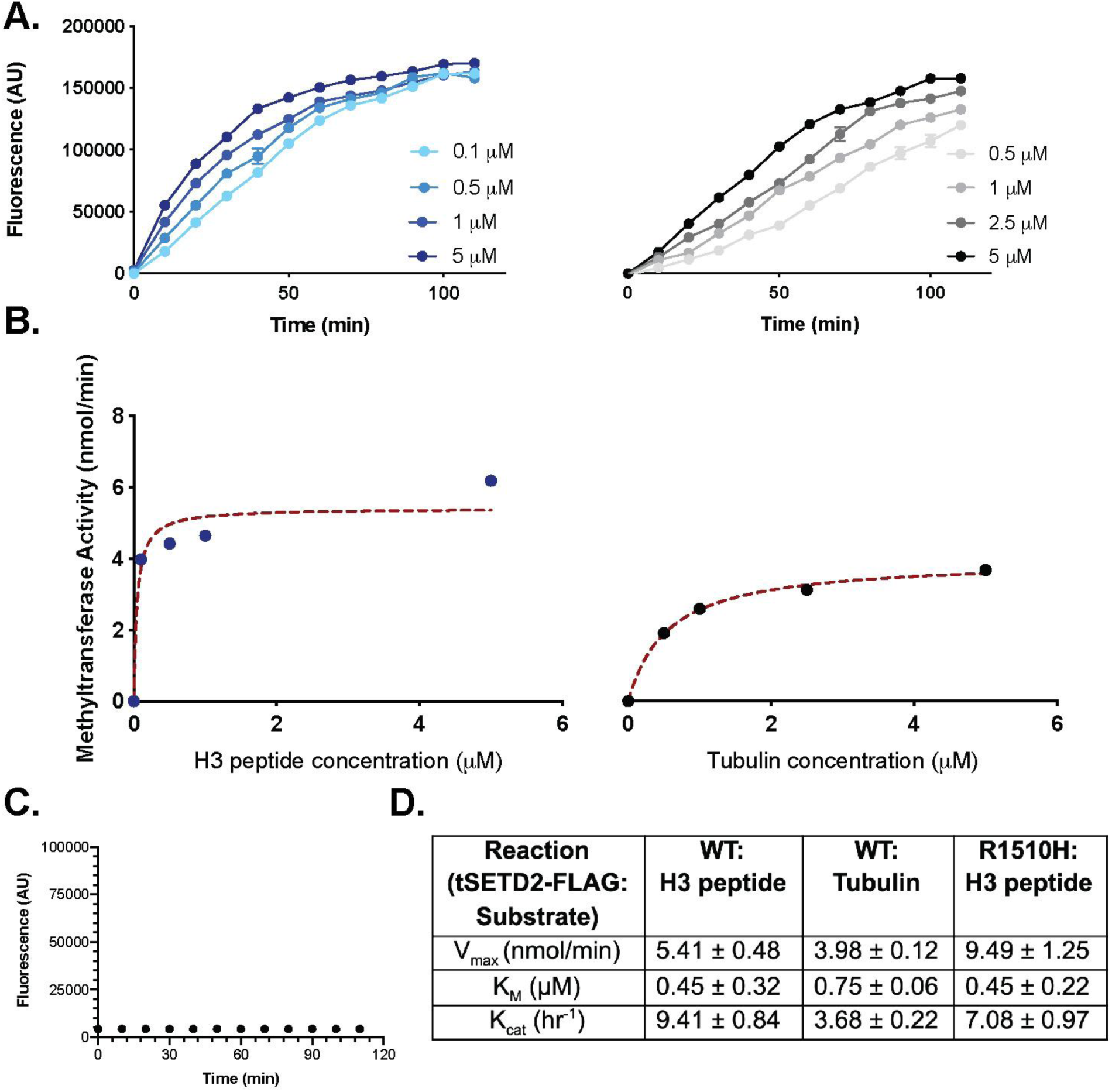
Kinetic analysis of tSETD2-FLAG methyltransferase activity. A) Fluorescence-based methyltransferase assay over time and with increasing amounts of substrate. tSETD2-FLAG was incubated with SAM and increasing amounts of either (left, blue) histone H3 peptide or (right, gray) porcine brain tubulin. B) Michaelis-Menten plots of substrate-dependent activity for (left, blue) histone H3 peptide or (right, gray) brain tubulin. Methyltransferase activity was calculated from the initial velocity values of the fluorescence plots in A) and a resorufin standard curve (RFU/concentration). Michaelis-Menten curve fits (dashed red lines) were obtained by least-squares method in Prism (8.1.1). C) Representative fluorescence trace of tSETD2-FLAG in the presence of SAM, but without substrate. Average values were background subtracted from reaction traces before kinetic analysis. D) Table showing kinetic values from methyltransferase assay for tSETD2-FLAG with histone H3 peptide or porcine brain tubulin.

**Figure S4:**
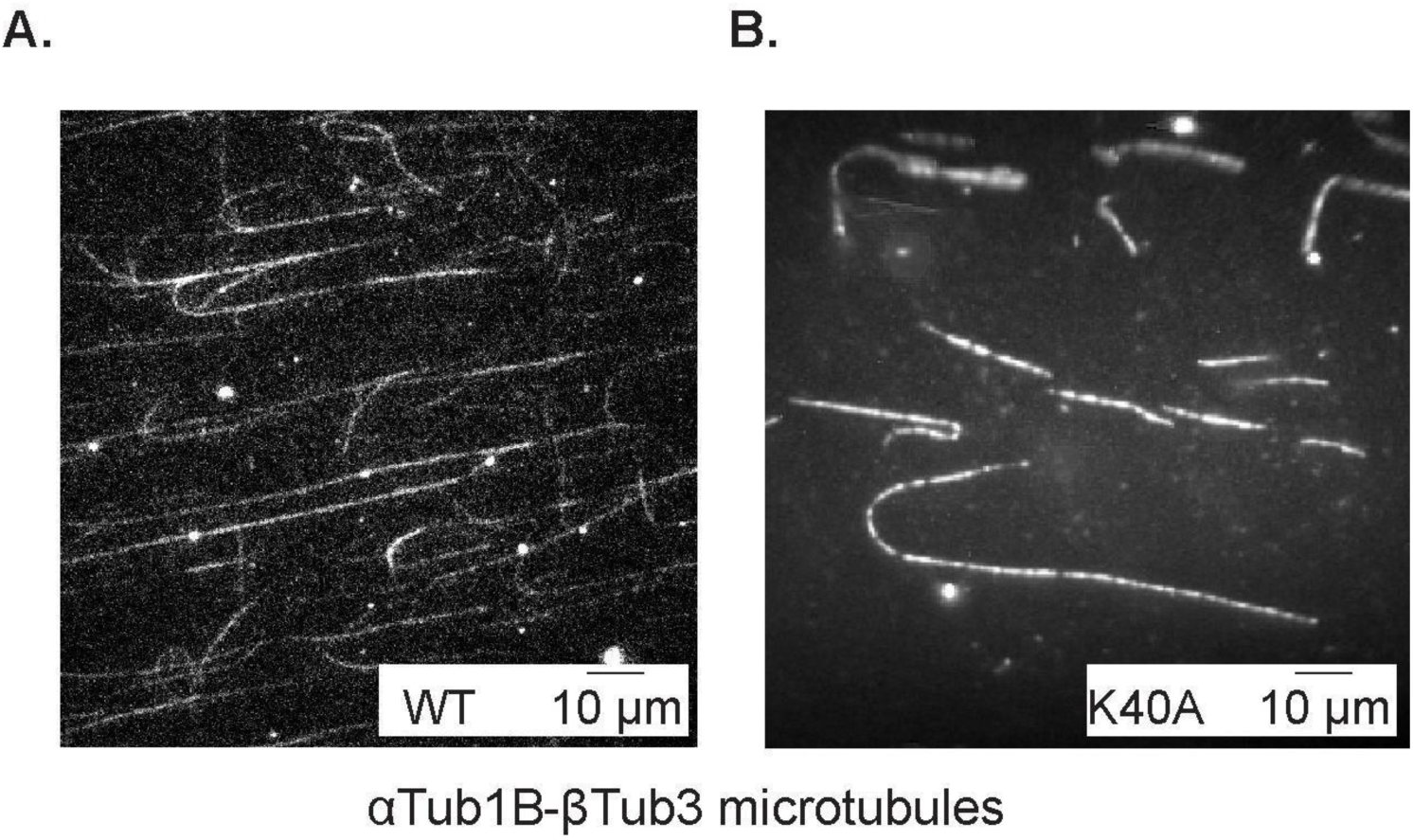
Polymerization of recombinant single-isotype tubulin into microtubules. Recombinant A) WT αβ-tubulin or B) mutant αβ-tubulin(αK40A) purified from insect cells was incubated with GMPCPP seeds to nucleate polymerization. To visualize growing microtubules, the reactions were spiked with 2% of 488- and biotin-labeled porcine brain tubulin. Shown is a representative field of viewed imaged by TIRF microscopy.

**Figure S5:** Mass spectra of all the samples *(to be deposited during final submission)*

**Figure S6:**
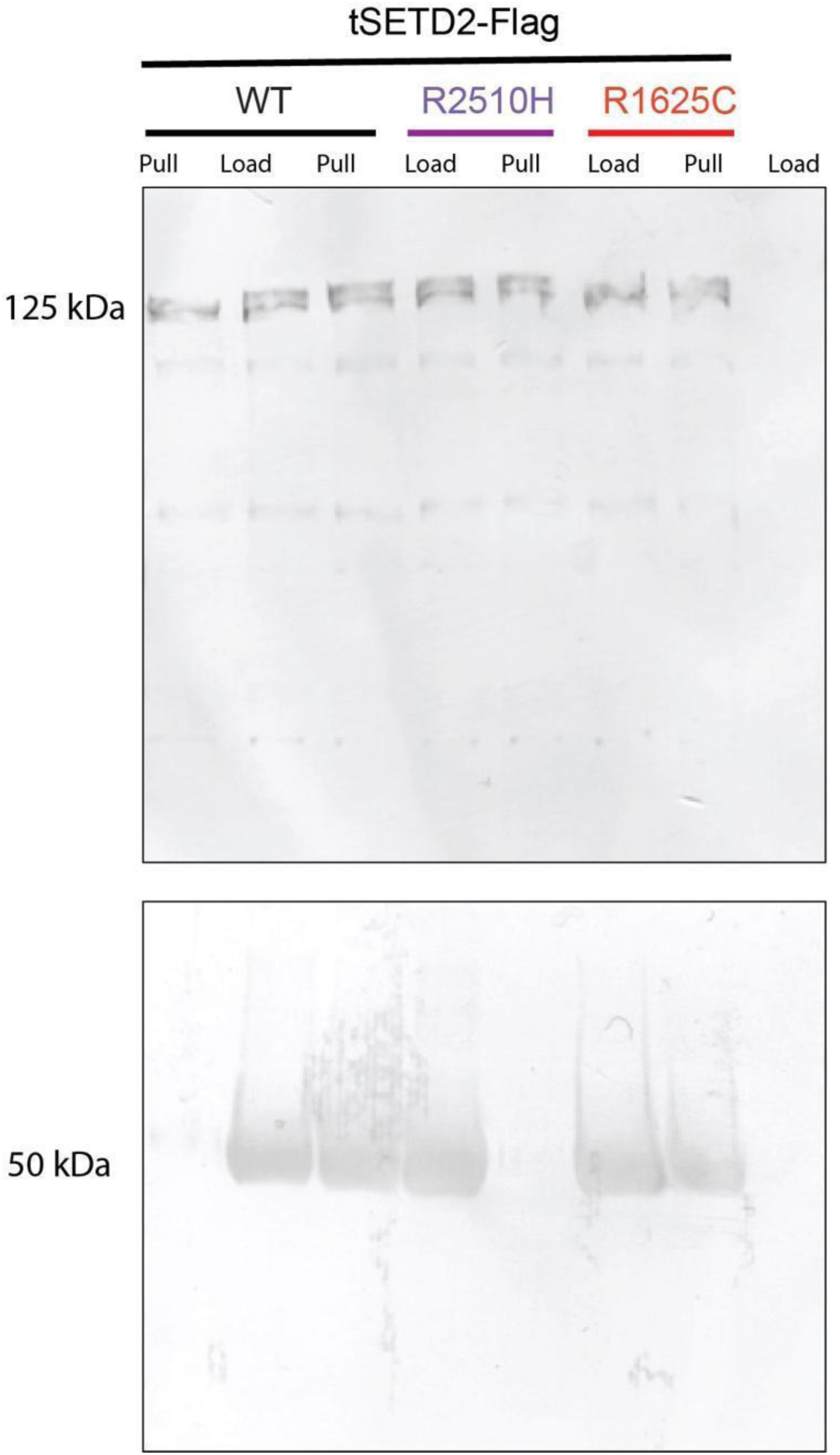
Related to Figure 4D. Uncropped view of western blots with top) anti-FLAG and bottom) anti-tubulin E7 antibodies.

**Figure S7:**
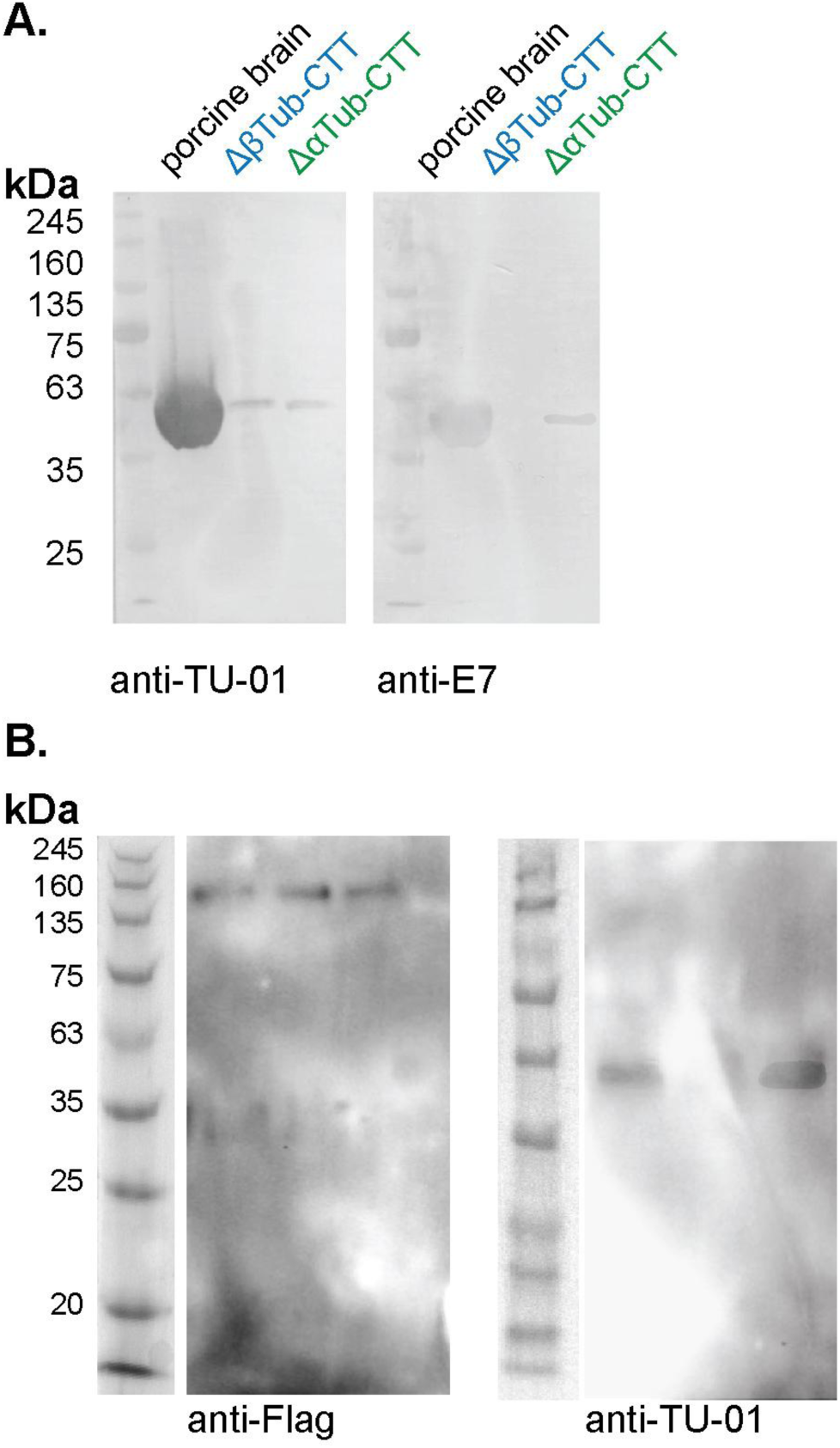
Related to Fig. 6. A) Uncropped view of western blots with antibodies to the left) α-tubulin N-terminus (TU-01) or right) β-tubulin CTT (E7). B) Uncropped view of IP western blots with antibodies to left) the FLAG tag and right) α-tubulin N-terminus (TU-01).

